# Impact of antibiotic perturbation on intestinal viral communities in mice

**DOI:** 10.1101/2021.10.28.466382

**Authors:** Jacqueline Moltzau Anderson, Tim Lachnit, Simone Lipinski, Maren Falk-Paulsen, Philip Rosenstiel

**Affiliations:** Institute of Clinical Molecular Biology, Christian-Albrechts University of Kiel, Kiel, Germany; Zoological Institute, Christian-Albrechts University of Kiel, Kiel, Germany

## Abstract

Viruses and bacteriophages have a strong impact on intestinal barrier function and the composition and functional properties of commensal bacterial communities. To improve our understanding of the role of the enteric intestinal virome, we longitudinally characterized the virome in fecal samples from wild-type (WT) C57BL/6J and knock-out (KO) NOD2 mice in response to an antibiotic perturbation. Sequencing of viral-like-particles (VLPs) demonstrated both a high diversity and high inter-individual variation of the murine gut virome composed of eukaryotic viruses and bacteriophages. Antibiotics also had a significant impact on the gut murine virome causing a delayed resilience independent of genotype. However, compositional shifts in the virome and bacteriome were highly correlated, suggesting that the loss of specific phages may contribute to a dysregulation of the bacterial community composition. Bacteriophage species may be playing an important role in either upregulating or downregulating the bacterial community, and restoring a healthy virome may therefore be a central goal of microbiota-targeted therapies.

## Introduction

Viruses are an integral part of the gut microbial community and are mostly comprised of bacteriophages [1, 2]. Previously, we demonstrated evidence of the role of NOD2 for controlling resilience of the intestinal microbiota (bacteriome and mycobiome), whereby the impaired recovery dynamics of the microbiota after antibiotic perturbation in NOD2-deficient mice is contributing to an inflammation-prone state of the intestinal mucosa. Such alteration in the capacity to restore a physiological equilibrium could be involved in the etiology of chronic inflammatory diseases and other intestinal disorders [3]. In favor of this hypothesis, it has been demonstrated in several human cohort studies that diversity and functional properties of the intestinal microbiota of Inflammatory Bowel Disease patients displays higher temporal fluctuation compared to healthy subjects, indicating a potential loss of control of the host. In contrast to the bacteriome, little is known about the intestinal virome in response to a specific pulse perturbation. Phages have been found to have various effects on the bacterial community, by impacting bacterial diversity in a community, stimulating evolutionary change, and providing selective advantages to their bacterial hosts [4]. Although it is obvious that shifts of bacterial taxa by specific antibiotics will directly cause secondary changes of the intestinal virome composition, it is likely that residing bacteriophages may exert an important level of control on the dynamics of bacterial community recovery by negative selection. Moreover, an enteric eukaryotic virus was shown to replace the beneficial function of the commensal bacteria in germ-free and antibiotic treated mice [5]. Thus, viruses can also play an important role in the regulation of intestinal homeostasis in response to antibiotic perturbations.

Viruses have also been shown to be extremely diverse, varying in their genetic material, genome sizes, life cycles, transmission routes, or persistence [6–9]. Humans are colonized by large populations of viruses consisting of viruses that infect eukaryotic cells (eukaryotic viruses) and those that infect bacteria (bacteriophages) [7, 10]. Human feces are estimated to contain at least 10^^9^ virus-like particles per gram [11], and although many of these viruses have been identified as bacteriophages, the majority remains unidentified [1, 2, 12]. Furthermore, host-genomes are also frequently composed of virus-derived genetic elements (retroviral elements and prophages) [7, 10, 13]. Metagenomic analyses of human gut viruses have also revealed extreme interpersonal diversity. This is in part likely due to the already considerable individual variation in the bacterial strains present in the gut, for which differences in phage predators are influenced [2, 14]. It is well established that phages can be highly selective for different bacteria, and as such, phage sensitivity (phage typing) has been used for decades as an effective means of differentiating between different bacterial strains [15, 16]. Rapid within-host viral evolution may also influence the large variability among individuals. In a long-term study investigating the viral community of an adult individual, Microviridae, a family of bacteriophages, was demonstrated to have high substitution rates, causing the sequence divergence values to be sufficient to distinguish new viral species by the conclusion of the study [11]. Moreover, individual virome compositions has been suggested to be relatively stable, with an estimated 80% of viral forms to be persistent throughout a 2.5 year-long study [11], with similar findings also observed in studies of shorter duration [1, 2].

Here, we demonstrate the effect of an antibiotic perturbation on the longitudinal variation of viral gut communities in C57BL/6J WT and NOD2 KO mice [3]. This community is largely uncharacterized, yet critical towards understanding its impact on the microbiome and health.

## Materials and Methods

### Animals

All animal experiments were approved by the local animal safety review board of the federal ministry of Schleswig Holstein and conducted according to national and international laws and policies (V 312-72241.121-33 [95-8/11]). All animals were housed in a mouse facility at the Christian Albrechts University of Kiel and experiments carried out as previously described [3].

Briefly, a single NOD2-deficient male mouse was crossed with a C57BL/6J female to obtain heterozygous offspring (F1), from which WT and NOD2 KO breeder pairs were generated (F2). Male offspring of the next two generations were then used and maintained in single cages under specific-pathogen free (SPF) conditions. At the onset of the study (Day 0), mice were approximately 52 weeks old. We treated C57BL/6J WT and NOD2 KO mice for two weeks with broad-spectrum antibiotics composed of ampicillin (1 g/L), vancomycin (500 mg/L), neomycin (1 g/L), and metronidazole (1 g/L) (Sigma Aldrich) [17], which were freshly prepared and administered ad libitum to the drinking water in light protected bottles. Fecal pellets were collected immediately throughout the 86 days of the study and stored at −80 °C until needed. Mice were monitored and weighed regularly and sacrificed at the conclusion of the study (Day 86). See Table S1 for more details on housing and samples.

### Virome Sample Processing

Two fecal pellets per sample were resuspended in 15 mL PBS buffer containing 0.01 M sodium sulfide and 10 mM EDTA for 30 min on ice. Samples were centrifuged twice at low speed (ThermoScientific Heraeus Multifuge 3SR) at 4°C for 30 min to remove bacteria and contaminating plant material. The resulting supernatants were sterile filtered and ultracentrifuged at 22,000 x rmp (Beckman SW41 rotor) at 4°C for 2 hrs. Viral pellets were then resuspended in 200 μL Tris buffer (50 mM Tris, 5 mM CaCl_2_, 1.5 mM MgCl_2_, pH 8.0), from which 5 μL sub-samples of isolated viruses were collected for morphological characterization by negative staining in 2% (w/v) aqueous uranyl acetate and visualized by transmission electron microscopy (TEM) (Technai Bio TWIN) at 80 kV with a magnification of 40,000-100,000. To the samples, 2 μL benzonase was added and incubated at 37°C for 2 hrs to remove remaining nucleic acid contamination.

To extract viral DNA and RNA, 22 μL of a 0.1 volume of 2M Tris-HCl (pH 8.5)/0.2 M EDTA, 10 μL of 0.5 M EDTA, and 268 μL of formamide were added to the sample and incubated at RT for 30 min. Subsequently, 1 μL of glycogen, and 1024 μL of ethanol were added, and samples were mixed gently and incubated overnight at RT. The next morning, samples were centrifuged at 12,000 x *g* at 4°C for 20 min, washed with 70% ethanol, and resuspended in 100 μL of TE buffer and 1 μL of mercaptoethanol, after which 10 μL of 10% SDS and 3 μL of Proteinase K were added and incubated for 20 min at 37°C and 15 min at 56°C. Then, 400 μL of DNA extraction buffer CTAB (100 mM Tris pH 8.0, 1.4 M NaCl, 20 mM EDTA, 2% CTAB) and 1 μL mercaptoethanol were added and samples were incubated at 56°C for 15 min. To the resulting supernatant, an equal volume of chloroform:isoamylalcohol (24:1) was added, and samples were centrifuged at 13,000 x *g* for 5 min. The supernatant was collected, to which 1 μL of glycogen, 10 μL mercaptoethanol, and a 0.7 volume of isopropanol were added and incubated overnight at −20°C. The next morning, samples were centrifuged at 13,000 x *g* at 4°C for 20 min, after which the supernatants were collected, washed with 500 μL of 70% ethanol, and stored at −80°C.

Following extraction of VLPs, ethanol was removed from samples and pellets were air-dried and resuspended in 20 μL of RNAse free filtered water. Amplification was performed using a modified Complete Whole Transcriptome Amplification Kit (WTA2) (Sigma-Aldrich) as described previously [18]. PCR products were then purified using the GenElute PCR Clean-Up Kit (Sigma-Aldrich). Samples were stored overnight at −20°C prior to library construction.

### Library Construction

Libraries were generated as described previously [18] using the NexteraXT kit (Illumina). After quantification, normalized pools of all samples were sequenced on an Illumina MiSeq using the 2 x 150bp sequencing kit (Illumina). This Whole Genome Shotgun project has been deposited with the links to BioProject accession number PRJNA434045 in the NCBI BioProject database (http://www.ebi.ac.uk/ena/data/view/PRJEB21817) with BioSample accession numbers SAMN08534315 through SAMN08534344.

### Viral community composition

Nextera XT adapters were removed and sequence reads were trimmed from Illumina paired-end reads (2×150 bp) using Trimmomatic V.0.36 [19]. Trimmed and quality controlled reads of all samples were cross assembled using SPAdes V.3.1.10 [20] to generate a reference viral metagenome. Contigs were screened for contamination by using blastn against the NCBI nucleotide database [21]. Contigs > 90% identity and > 50% of length were removed if no viral hallmark gene could be detected within the sequence. Contigs with a minimum length of 1,000 bp and a minimum total read coverage of 10 were selected and analyzed using blastx against the UniProt viral database including 5,571,160 viral sequences with an e-value cut off at 10^-5^ [22]. Finally, the reference viral metagenome was further classified by VirSorter2 [23] and contig annotation tool CAT [24]. Moreover, all contigs were submitted to Rapid Annotation using Subsystem Technology (RAST) to identify additional viral hallmark genes. Moreover, VirHostMatcherNet [25] and CAT [24] were used to predict virus-prokaryote interaction. Contigs classified as virus by VirSorter2, CAT or RAST were used as OTUs representing the mice viral community. Reads from each sample were then mapped separately against representative mice viral OTUs using the computer software Bowtie2 [26] and SAM tools [27]. The normalized coverage of each OTU was used as a proxy for the relative abundance of each virus per sample [28].

Viral community composition was analyzed using the computer software PRIMER V.7 [29–31], and abundance data was standardized and log+1 transformed. Estimation of similarity between all samples was calculated by Bray-Curtis similarity and non-metric multidimensional scaling analysis (MDS), and pairwise comparison of viral community composition between different treatment groups and time points was analyzed using a similarity test (ANOSIM global test) [31]. SIMPER analysis was used to detect the most important viral OTUs that contribute to observed difference in community composition. These preselected OTUs were further analysed by one-factor ANOVA followed by Turkey’s honest significant differences (HDS) test using the computer software (SPSS).

### Relationship between viral and bacterial community

To investigate the variability in the viral community that could be explained by the bacterial community composition, or vice versa, RELATE analysis [30] in the computer software PRIMER V.7 [29, 31] was used. The analysis was based on the relative abundances of viral and bacterial OTUs. Raw bacterial FASTQ reads were obtained from our previous study [3] from EBI’s ENA under the Accession Number PRJEB21817 (http://www.ebi.ac.uk/ena/data/view/PRJEB21817). Viral and bacterial community datasets were standardized and log(x+1) transformed. To investigate the variability of the bacterial community composition that could be explained by the viral community, we fitted the 29 most abundant viral OTUs with a minimum length of 10,000 bp to the relative abundance of bacterial OTUs using distance-based redundancy modeling (DISLM) with adjusted R^2^ selection criteria and forward selection procedure. Results were visualized with distance-based redundancy analysis (dbRDA) [32, 33].

## Results

### Presence of virus-like particles

The presence of virus-like particles in fecal samples was observed by transmission electron microscopy, which revealed morphologically distinct isolates (Fig. 1). Numerous diverse bacteriophages were present (Fig. 1a-c), and were distinguished by the structure of a head, or capsid, and in some cases a tail, although other phage morphologies exist beyond this structure (*i.e*. without a tail). The *Myoviridae* family morphology was present, with an icosahedral (20 sides) head and a rigid tail (Fig. 1c). The structure of a phage lambda (λ) displaying the *Siphoviridae* family morphology, which commonly infects *E. coli*, was also identified (Fig. 1a, b). The protein head of the capsid is icosahedral (Fig. 1b) and elongated (Fig. 1a), containing the nucleic acid. The head is joined to a tail possessing a long thin tail fiber at its end (for host recognition). The tails are composed of a hollow tube, through which the nucleic acid passes into the host during infection. Virus-like particles with morphological similarity to eukaryotic viruses, e.g. the *Peste des Petits Ruminants* (PPR) virus or the *Murine Mammary Tumor* virus (MMTV) were also observed (Fig. 1d-f).

**Figure 1.**
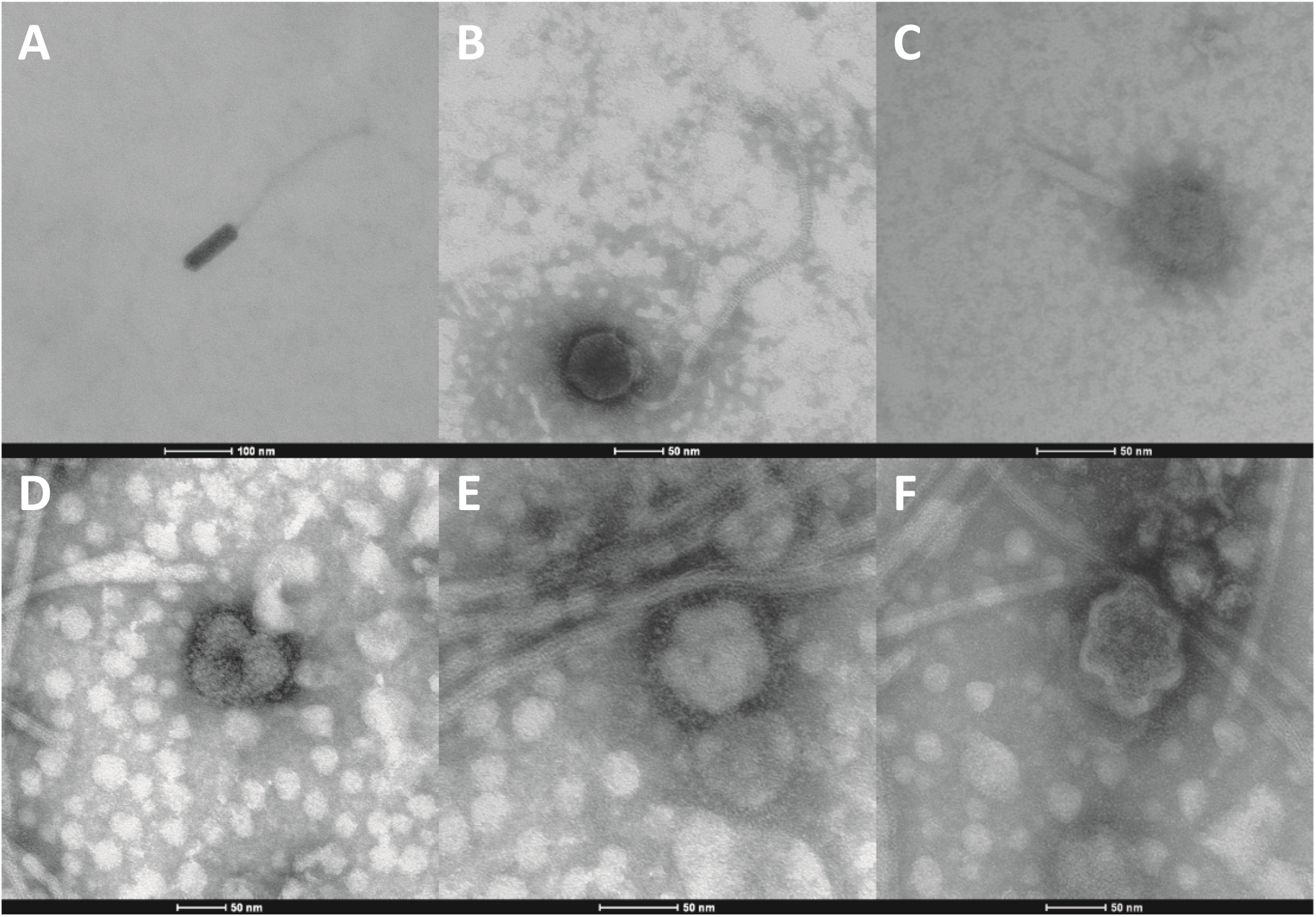
Transmission electron micrographs (TEM) of purified virus-like particles from murine feces negatively stained with 2% aqueous uranyl acetate. **(A-C)** Bacteriophages from the family Siphoviridae and Myoviridae, respectively, and **(D-F)** Eukaryotic viruses with morphological similarity to PPR virus or MMTV virus.

### Reference murine fecal virome composition

Sequence reads of all murine fecal viruses were assembled into 1,094,102 contigs. For our reference virome, we selected 4,767 contigs that were longer than 1,000 bp and had a coverage higher than 10. The reference virome had an average sequence length of 3,358 and a coverage of 80. Of these contigs, 48% were assigned to known viral sequences using the Uniprot viral database. To reduce the impact of false positives we focused our viral community analysis only on contigs that were assigned as viral sequences based on VirSorter2, CAT and RAST annotation. This subset consisted of 614 contigs composed of approximately 94% dsDNA viruses, 5% ssDNA viruses, 1% RNA viruses.

The viral community was predominantly composed of bacteriophages, consisting primarily of the order *Caudovirales* (71% of the viral contigs). Approximately 16% of the viral contigs were predicted by VirSorter2 as the eukaryotic viruses *Lavidaviridae* and *NCLDV*. To reduce the false positive detection of eukaryotic viruses, these contigs were compared by blastn and blastx to NCBI’s non-redundant protein database, most of which were found to have a high sequence similarity to prokaryotes rather than eukaryotes. A few viral sequences were identified in murine feces that infect eukaryotes. One ssRNA virus of the family *Retroviridae* was found showing high sequence similarity on a nucleotide level to *Murine leukemia virus* (contig 3291).

A dsRNA virus *Hordeum vulgare alphaendornavirus* (contig 811, 1176 and 3773) of the family *Endornaviridae* was found which is known to infect barley. Other potential plant associated viruses that could be identified in this study were ssDNA viruses of the family *Genomoviridae* with high sequence similarity to *Gemycircularvirus* (contig 4238 and contig 2208).

### Delayed resilience in viral gut community composition post antibiotic perturbation

Multidimensional scaling analysis of the viral community based on viral OTU level demonstrated a clear clustering based on day (Fig. 2). Samples at Day 0, prior to treatment, clustered together and communities underwent significant changes shifting after 14 days of antibiotic treatment (ANOSIM global test for test differences between time points: R statistics = 0.443, *P* = 0.001). Differences between the genotypes could not be detected at any time point (ANOSIM global test for differences between time points: R statistic = −0.03, *P* = 0.753). Antibiotic treatment significantly changed the viral community composition ANOSIM Pairwise Test Supplementary Information S2). 20% of the average dissimilarity between Day 0 and Day 14 was explained by the higher relative abundances of 4 phages infecting *Gammaproteobacteria* and a reduction of two phages infecting *Bacteroides* bacteria after antibiotic treatment (SIMPER analysis; Supplementary Information S3). We could confirm by ANOVA that *Escherichia* phages (contigs 14, 32, 186, and 3817) increased, whereas phages predicted to infect *Bacteroidetes*, such as Phage *apr34* (contig 52), and *Microvirus* (contigs 996) were reduced after antibiotic treatment (n = 6, *P* < 0.01, one-way ANOVA, Tukey’s HDS).

**Figure 2.**
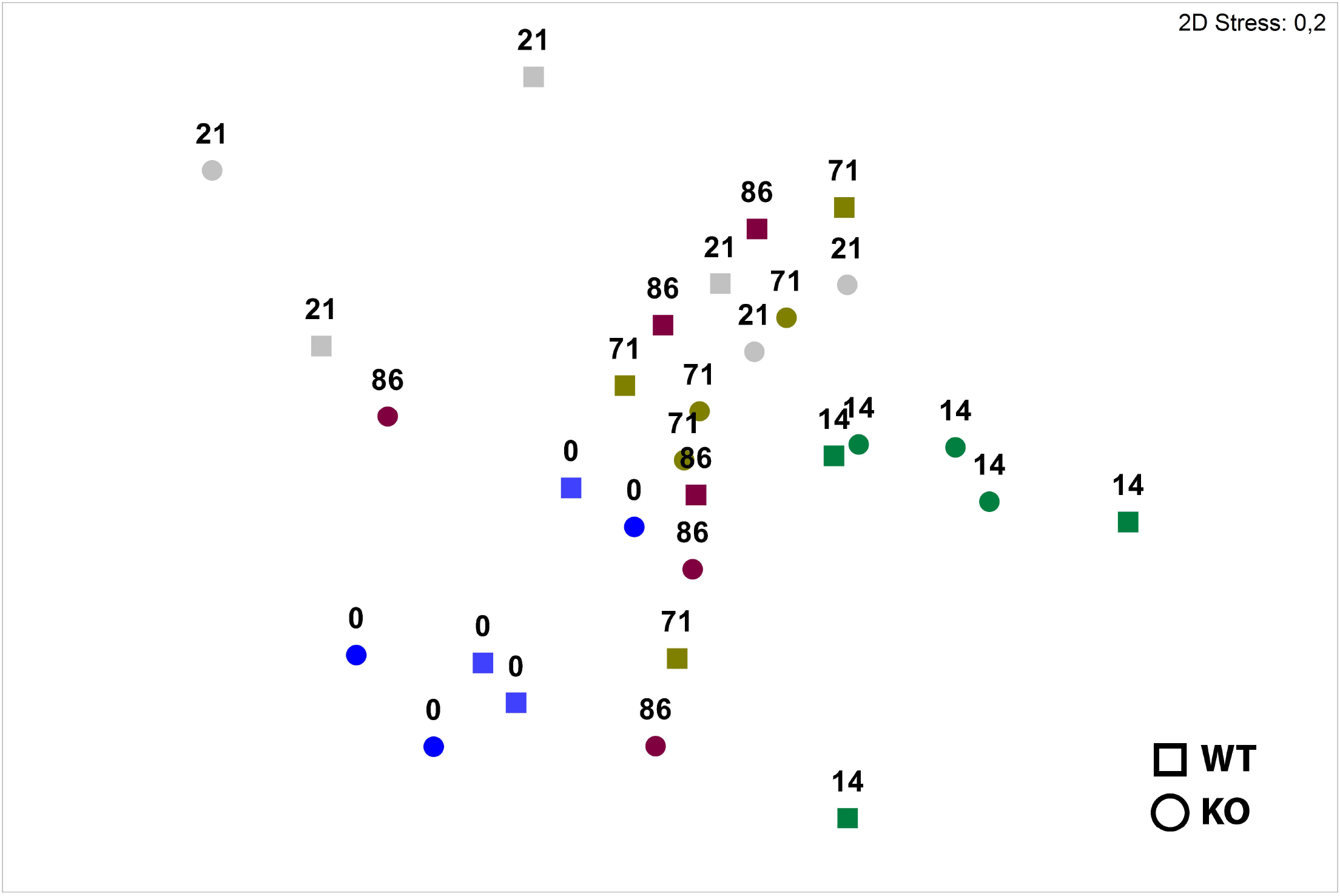
Non-metric mutidimentional scaling (NMDS) analysis of the murine fecal viral community composition of NOD2 KO (triangles) and C57BL/6J WT (circles). Analysis based on the Bray-Curtis similarity index of the relative abundance of viral OTUs at species level across time (Day 0 = pre-treatment, Days 14-86 = post-treatment).

The community compositional trajectory shifted towards recovery with increasing time by clustering more closely with Day 0 (developing towards a community composition similar to prior antibiotic treatment) (Fig. 2). However, viral community composition at Day 86 did not fully recover and remained significantly different from Day 0 with an average dissimilarity of 86.64 (ANOSIM Pairwise Test for differences between day 0 and day 86: R statistics = 0.581, *P* = 0.002). Of this dissimilarity, 20% could be explained by as few as 6 viral OTUs, of which *Bacteroidetes* infecting phages *Microvirus* (contig 935) and Phage apr34 (contig 52) were highly reduced after the antibiotic perturbation and could not be detected at the end of the study (Day 86) (Table S4). Moreover, comparing viral diversity pre-antibiotic treatment compared to post-treatment, Day 0 (pre-treatment) had a significantly higher viral diversity in both total species (n = 6, F = 13.525, *P* < 0.001, one-way ANOVA, Tukey’s HDS) and species richness (Margalef) (n = 6, F = 13.527, *P* < 0.001, one-way ANOVA, Tukey’s HDS) (Fig. 3).

**Figure 3.**
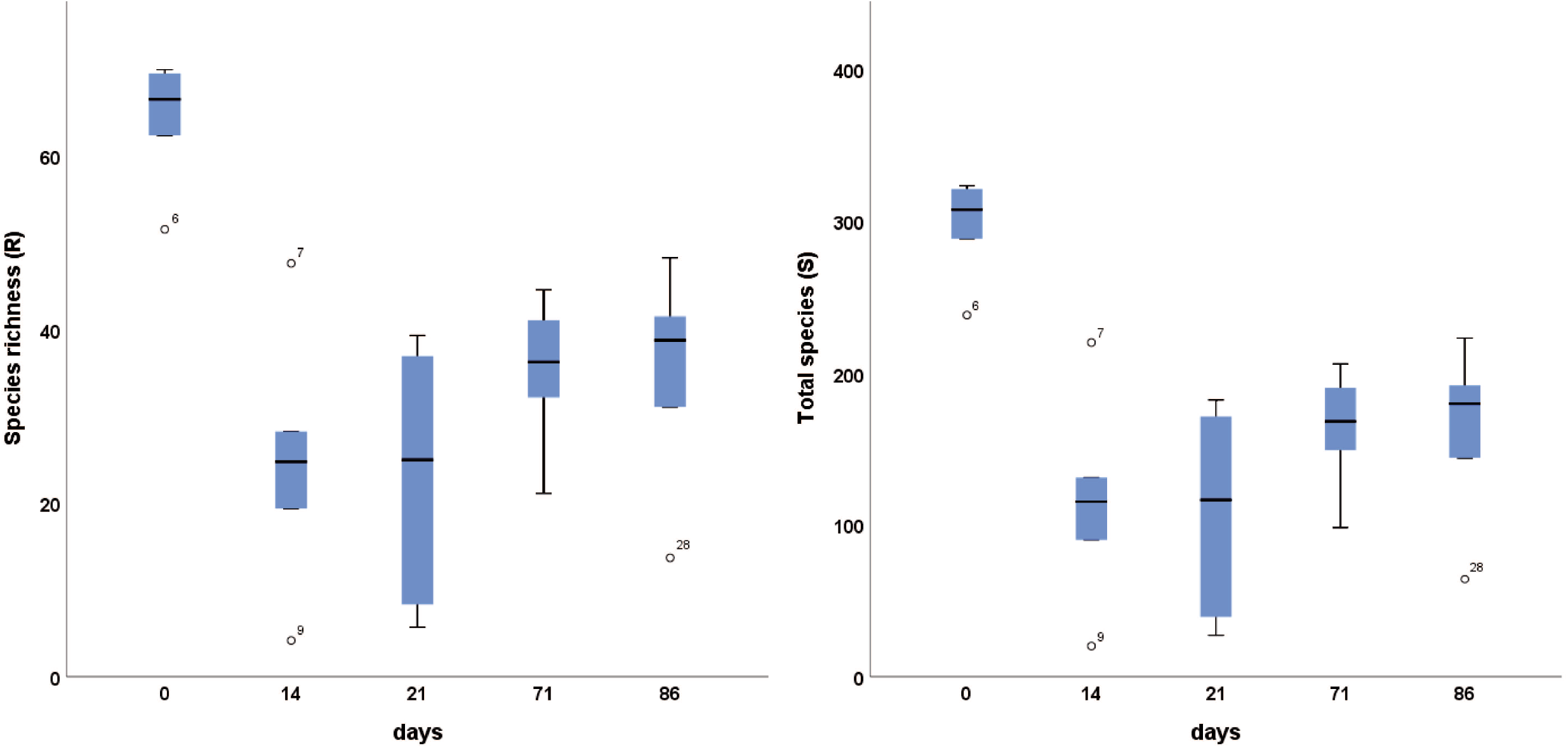
**(A)** Viral community composition of the reference virome from sequence reads of all murine fecal samples. Contigs were assigned to known viral sequences by the UniProt viral database. **(B)** Prokaryotic viral community composition in KO and WT mice across time (Day 0 = pre-treatment, Day 14 = antibiotic treatment, Days 21-86 = post-treatment). Bacteriophages were grouped according to their blastx hits in the UniProt viral sequence database. Roman numbers represent individual mice.

### High inter-individual variation of prokaryote viral community composition

Fecal bacteriophages were diverse and variable in their relative abundance between different individual mice. Prior to antibiotics (Day 0) dominant phages were *Microviridae*, *CrASSphage*, Phage *apr34* and other *Bacteroidetes* and *Firmicutes* infecting phages (Figure 4). Antibiotic perturbation strongly affected phage composition shifting to an *Escherichia* phage dominated system. Within one week of antibiotic cessation the viral community composition demonstrated huge variability. No clear pattern of recovery over time or between the genotypes could be observed, with high inter-individual variation of viral community composition present throughout (Fig. 4). The analysis on an OTU level indicated shifts in *Gammaproteobacteria*, *Firmicutes*, and *Bacteroidetes* phages. To determine whether these shifts in the phage population were significant, we used VirHostMatcherNet and CAT taxonomy for bacterial host prediction. All phages were grouped based on their bacterial host prediction at a higher phylogenetic level (*i.e. Gammaproteobacteria*, *Firmicutes*, and *Bacteroidetes*) (Fig. 4). Prior to the antibiotic treatment, the phage population was dominated by equal portions of *Bacteroidetes* and *Firmicutes* phages. Large changes within the community composition occurred during antibiotic treatment (Day 14), which was distinct from pre-treatment at Day 0 (Figures 2 and 4). During this time, the phage community was dominated by *Gammaproteobacteria* phages (Figure 5).

**Figure 4.**
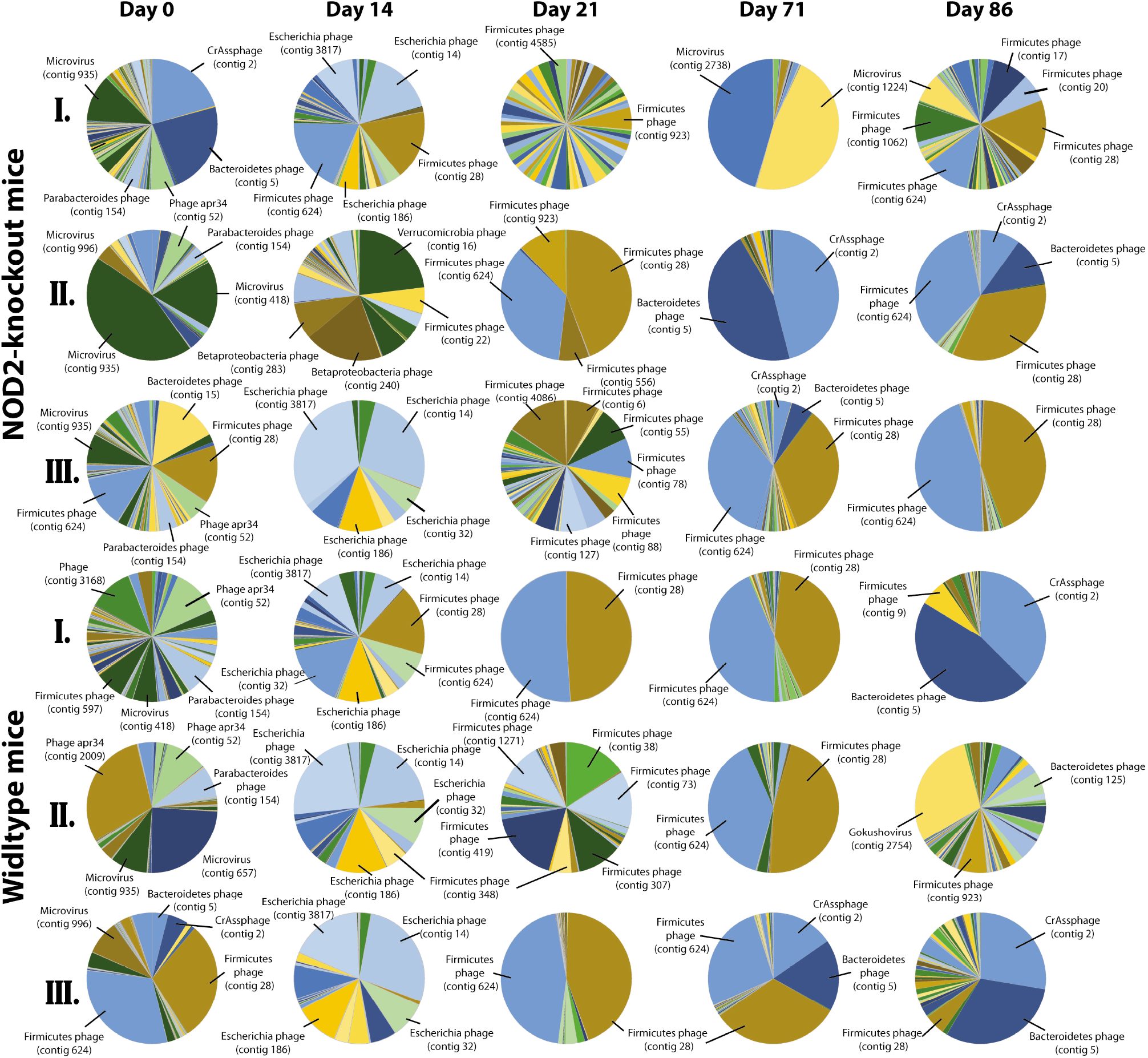
Relative abundance (%) of **(A)** *Bacteroidetes*, *Firmicutes*, and *Gammaproteobacteria* phages, compared to the relative abundance (%) of the corresponding bacteria. Phages were grouped based on their classification by blastx against the UniProt viral database. Time is represented in days (Day 0 = pre-treatment, Day 14 = antibiotic treatment, Days 21-86 = post-treatment). (Error bars = +/− 2 SE, n = 3).

**Figure 5.**
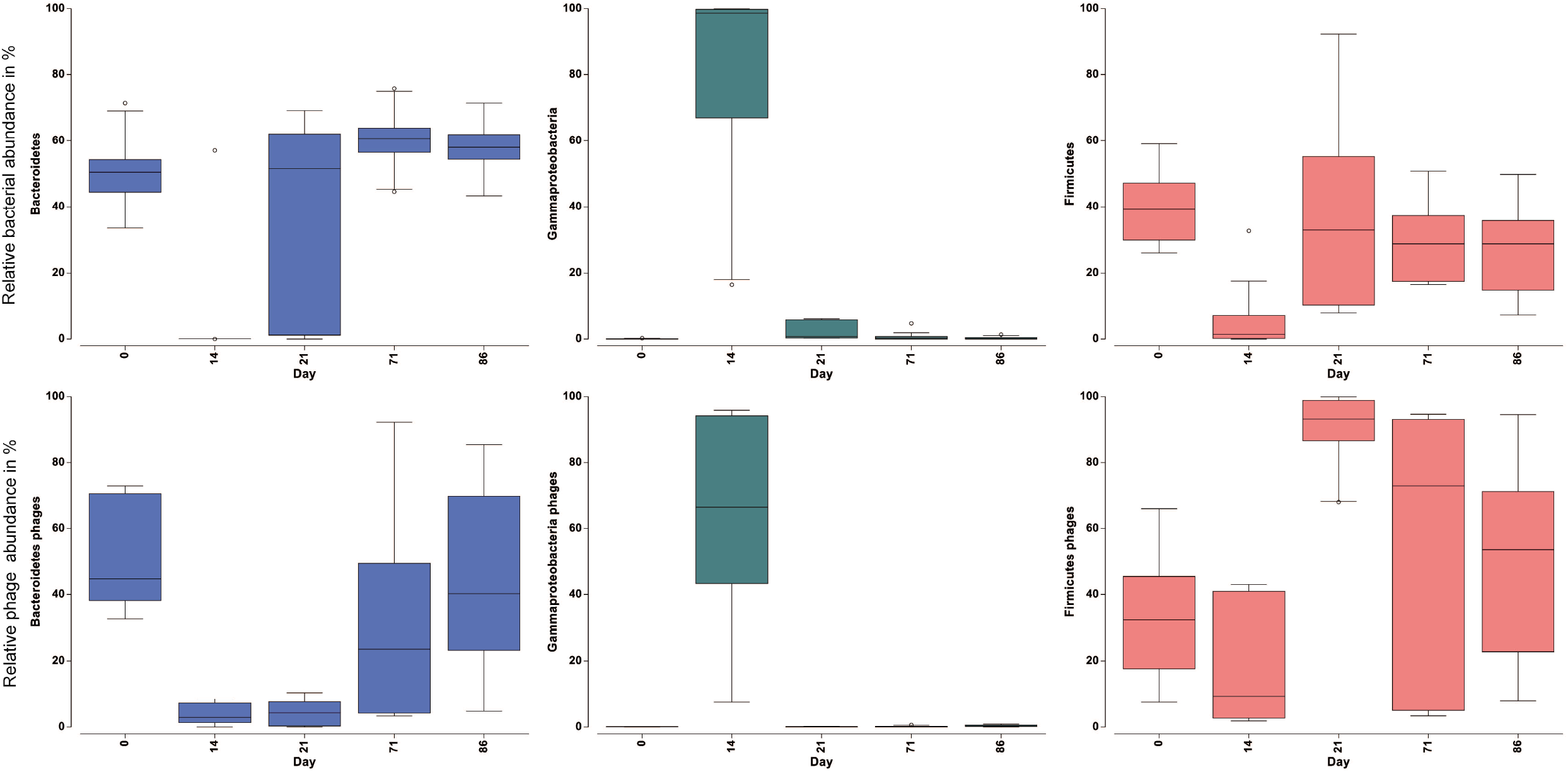
Constrained dbRDA plot of the murine fecal viral community composition of KO (triangles) and WT (circles) across time fitted to the fecal bacterial community composition in DistLM sequential tests with R^2^ selection criteria and forward selection procedure. Lengths of vector overlays indicate the relative influences of related predictor variables. Time is represented in days (Day 0 = pre-treatment, Day 14 = antibiotic treatment, Days 21-86 = post-treatment).

After antibiotic perturbation, *Bacteroidetes* phages were significantly reduced from 44.8 % to 5.3 % at day 21 (n = 6, F = 3.475, *P* = 0.025, one-way ANOVA, Tukey’s HDS). The relative abundances of Firmicutes phages were only slightly affected by antibiotic perturbation from 40 % to 17.8 % at the end of antibiotic treatment (day 14), not significantly affected compared to pre-antibiotic treatment. During the resilience period (post Day 14), the relative abundance of *Gammaproteobacteria* phages was reduced, while *Firmicutes* phages recovered rapidly and reached higher relative abundance compared to pre-antibiotic perturbation at day 21 (n = 6, F = 5.78, *P* = 0.013, one-way ANOVA, Tukey’s HDS). This was in contradiction to observed shifts in the bacterial communities, which featured higher relative abundances of *Bacteroidetes* bacteria compared to *Firmicutes* bacteria at Day 71 (n = 6, F = 20.4, *P* = 0.001, one-way ANOVA, Tukey’s HDS) and Day 86 (n = 6, F = 18.3, *P* = 0.002, one-way ANOVA, Tukey’s HDS). In contrast to *Firmicutes* phages, recovery of *Bacteroidetes* phages was delayed and only detectable from day 71. Interestingly, *Bacteriodetes* and *Firmicutes* bacteria were differently affected by antibiotic perturbation showing an almost total eradication of *Bacteroidetes* after 14 days of antibiotic treatment, while *Firmicutes* bacterial population still reached 7% relative abundance at day 14.

Changes in the compositional shifts of both bacterial and viral communities were correlated, where changes in the bacterial community composition (based on an OTU level) occurred in a similar direction and magnitude as the compositional shifts of the viral community (RELATE, Rho = 0.416, *P* = 0.001, 999 permutations). Time after the antibiotic perturbation (resilience period) was the main factor in both groups. Using distance-based modelling (DISLM) of the 29 most important phages, the relative abundance of *Escherichia phage* (contig 14) was found to be higher from the antibiotic perturbation and explained 25.5% of the variation in the bacterial community (Fig.6). Together with four other bacteriophages, the model could explain up to 49% of the bacterial variation (Fig. 6 and Table S5).

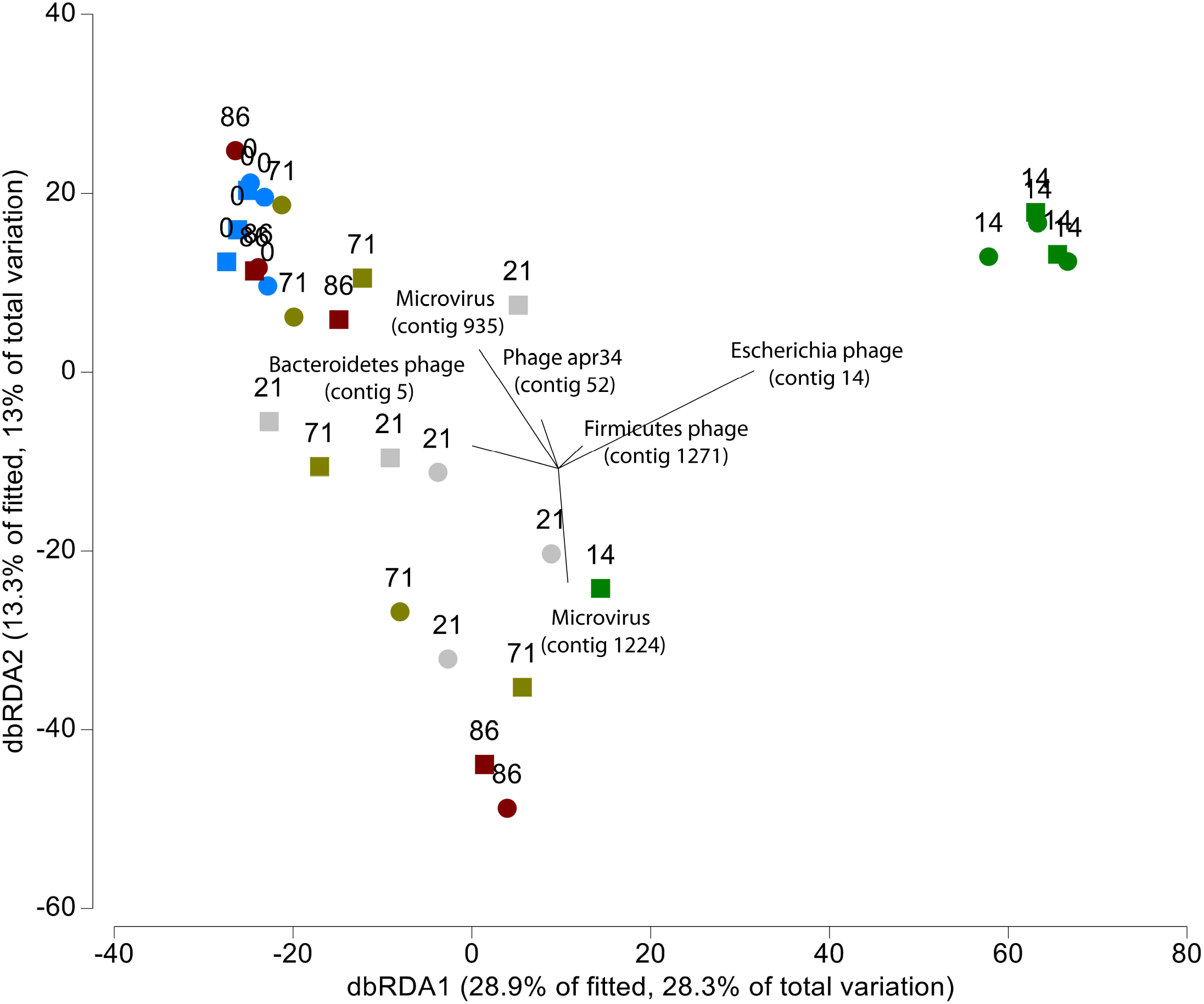

## Discussion

Investigations of the gut microbiota have largely ignored the virome, and the inclusion of RNA viruses in these studies has been further overlooked. To our knowledge, resilience properties of the virome post an antibiotic perturbation have never been explored. Here, we demonstrated that the murine gut virome is morphologically and genetically diverse, including viruses infecting the host (eukaryotes, *i.e. Retroviridae*, *Murine leukemia virus*, *etc*.), viruses infecting prokaryotes (bacteriophages, *i.e*. *Caudovirales*), and viruses infecting neither of them (plant viruses, *i.e. Hordeum vulgare alphaendornavirus*).

The murine gut virome shares several characteristics to the human gut virome. Firstly, the murine gut virome was highly variable among individuals. This high inter-individual viral diversity has also been previously reported in the human gut virome [1, 34]. Furthermore, similarly to the human gut virome, the murine gut virome was established to contain a large diversity of primarily bacteriophages, in addition to a much lower diversity of eukaryotic viruses [1, 2, 11]. Additionally, consistent with previous reports, the most abundant viral taxa identified were bacteriophages of the order Caudovirales (*i.e. Cellulophaga* phage) and the family Microviridae (*i.e. Parabacteroides* phage) [1, 2, 34].

RNA viruses associated with murine feces were also identified, and ultimately, only these RNA viruses could be verified as eukaryotic viruses. Of these eukaryotic viruses, some were identified as having a plant host (*i.e. Hordeum vulgare alphaendornavirus*). Plant viral sequences likely reflect the omnivorous diet of these mice, and we speculate that diet plays a significant role in the acquisition of the gut eukaryotic virome. Interestingly, the detection of these viruses was predominantly from Day 14, which had the lowest phage diversity. We further speculate that these plant viruses are always present, but could not be detected as a result of the greater phage community. Most studies to date have not reported the presence of large eukaryotic viruses as a result of filtering methods during isolation and extraction [18]. For instance, although filtering methods efficiently remove bacteria, filters have also been shown to remove more than 99% of *Mimivirus* and 90% of herpes viruses [18]. It is possible that these viruses are present more often than originally considered, though it remains to be seen what role they may play within the host. It should also be mentioned that there is an urgent need to develop improved references for characterizing the virome, as evidenced by the large percentage of sequences that were unclassified in the assembled reference, yet originated from viral fractions. The resulting sequencing catalog generated from this study is composed of nearly full genomes of high quality, serving as an important reference for future virome studies.

Prior to antibiotic perturbation, the phage population was dominated by equal proportions of *Bacteroidetes* phages and *Firmicutes* phages, reflecting the bacterial community which was dominated by *Bacteroidetes* and *Firmicutes*. During the antibiotic perturbation, a significant change occurred on the viral community composition. Although antibiotics do not directly target viruses, our results demonstrated significant changes in the bacterial community, which in turn largely impacted the gut bacteriophage community. During this time, *Bacteroidetes* phages were largely reduced and replaced by *Gammaproteobacteria* phages. It is possible that the outgrowth of *E.coli/Shigella* during antibiotic treatment led to the bloom of their respective bacteriophage *(i.e. Escherichia shigella* phage) as expected in Lotka-Volterra/Kill-the-Winner dynamics [14, 35]. Interestingly, numerous studies have demonstrated a significant association between the NOD2 risk allele and the increase in relative abundance of *Enterobacteriaceae* [36–38]. However, contrary to a previous study in humans, no unique changes occurred in the bacteriophage community specific to the NOD2 KO [34]. After the antibiotic perturbation, compositional shifts in the murine bacterial and viral communities were significantly correlated, where changes in the communities occurred in a similar direction and magnitude. *Gammaproteobacteria* phages were reduced, whereas an increase in Firmicutes phages occurred and remained until the end of the study. These changes in the phage community were also reflected in the bacterial community through an increase in the phyla *Firmicutes*. On the other hand, while *Bacteroidetes* phages did increase after Day 21, their relative abundance remained low compared to Day 0. This was in contradiction to observed shifts in the bacterial community, which featured higher relative abundances of *Bacteroidetes* bacteria compared to Firmicutes bacteria at Days 71 and 86. It is possible that the loss of *Croceibacter* phage, which disappeared during the antibiotic perturbation, may have been a regulator of *Bacteroidetes* bacteria, further revealing that virus-bacteria community dynamics of the gut are complex. Moreover, the identification of *Firmicutes* and *Bacteroidetes* phages associated with their bacterial phyla (*i.e. Firmicutes* and *Bacteroidetes)* both prior to the perturbation and at the conclusion of the study, suggests an important role in the virus-bacterial dynamics of these communities in maintaining host health. Nonetheless, ultimately the viral community had an impaired recovery, though the community appeared to be re-approaching a structure similar to Day 0 (pre-treatment).

Taken together, the antibiotic perturbation caused a delayed recovery in the gut virome independent of genotype. The perturbation led to substantial shifts in the murine gut viral community, further emphasizing the beneficial and detrimental effects viruses can have in response to environmental and host factors. In particular, the results presented here indicated that bacteriophage species may be playing an important role in either upregulating or downregulating the bacterial community, and their loss might contribute to a disturbed microbiome. Restoring a healthy virome may therefore be a central goal of microbiota-targeted therapies, which could be a disruptive approach in a variety of intestinal disorders, from IBD to colorectal cancer, and highlights the importance of better understanding factors contributing to resilience.

## Acknowledgments

The authors thank Melanie Vollstedt for their technical assistance. This work was supported by the IMPRS of the Max-Planck-Institute for Evolutionary Biology, the DFG Excellence Cluster 2167 Precision Medicine in Chronic Inflammation (TI-1 to P.R.), and DFG collaborative research center (CRC) 1182 Origin and Function of Metaorganisms, C2 (P.R.) and C4.2 (T.L.). Sequencing was supported by the DFG Sequencing Hub CCGA at CAU Kiel (P.R.).

---

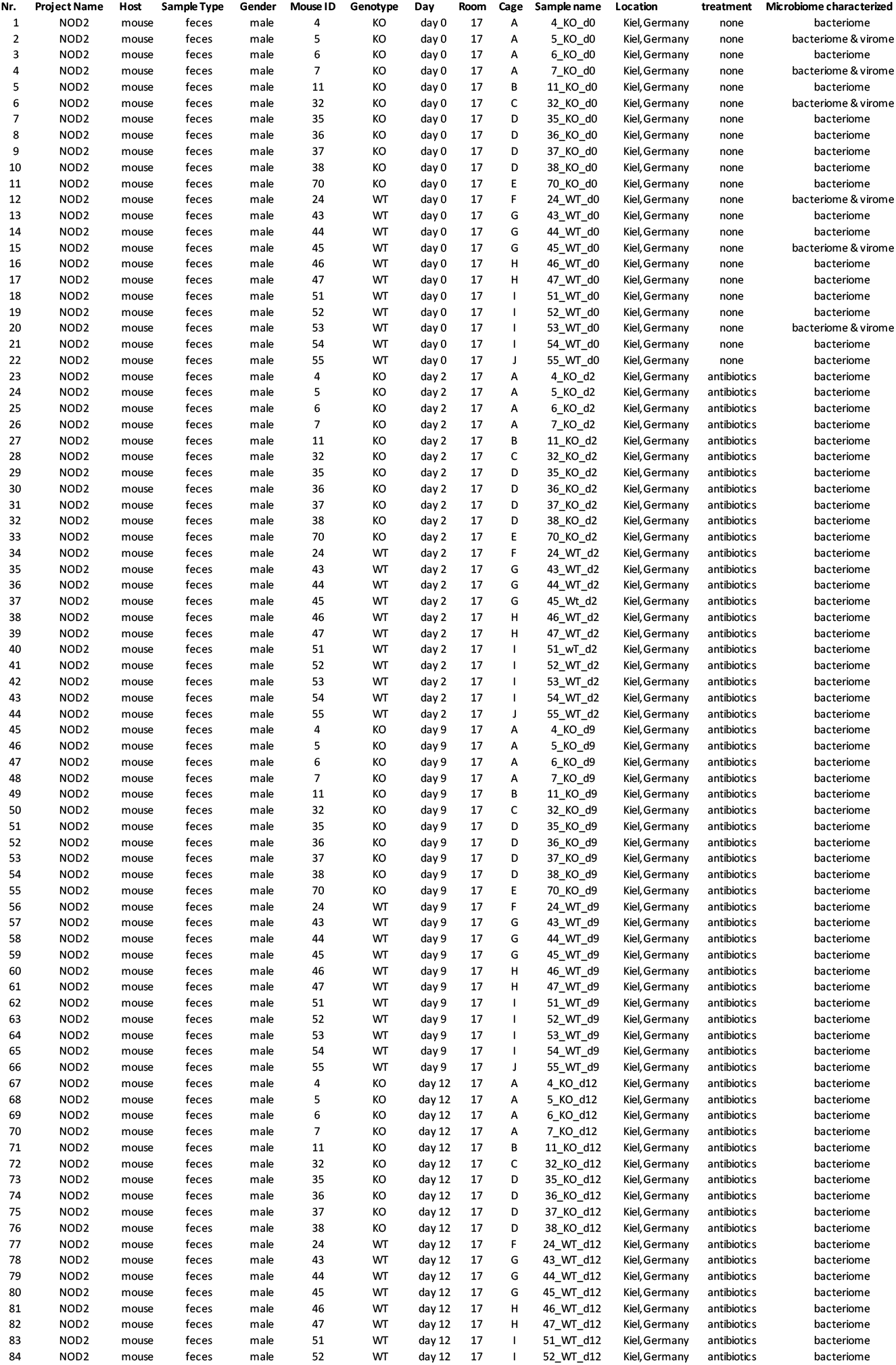
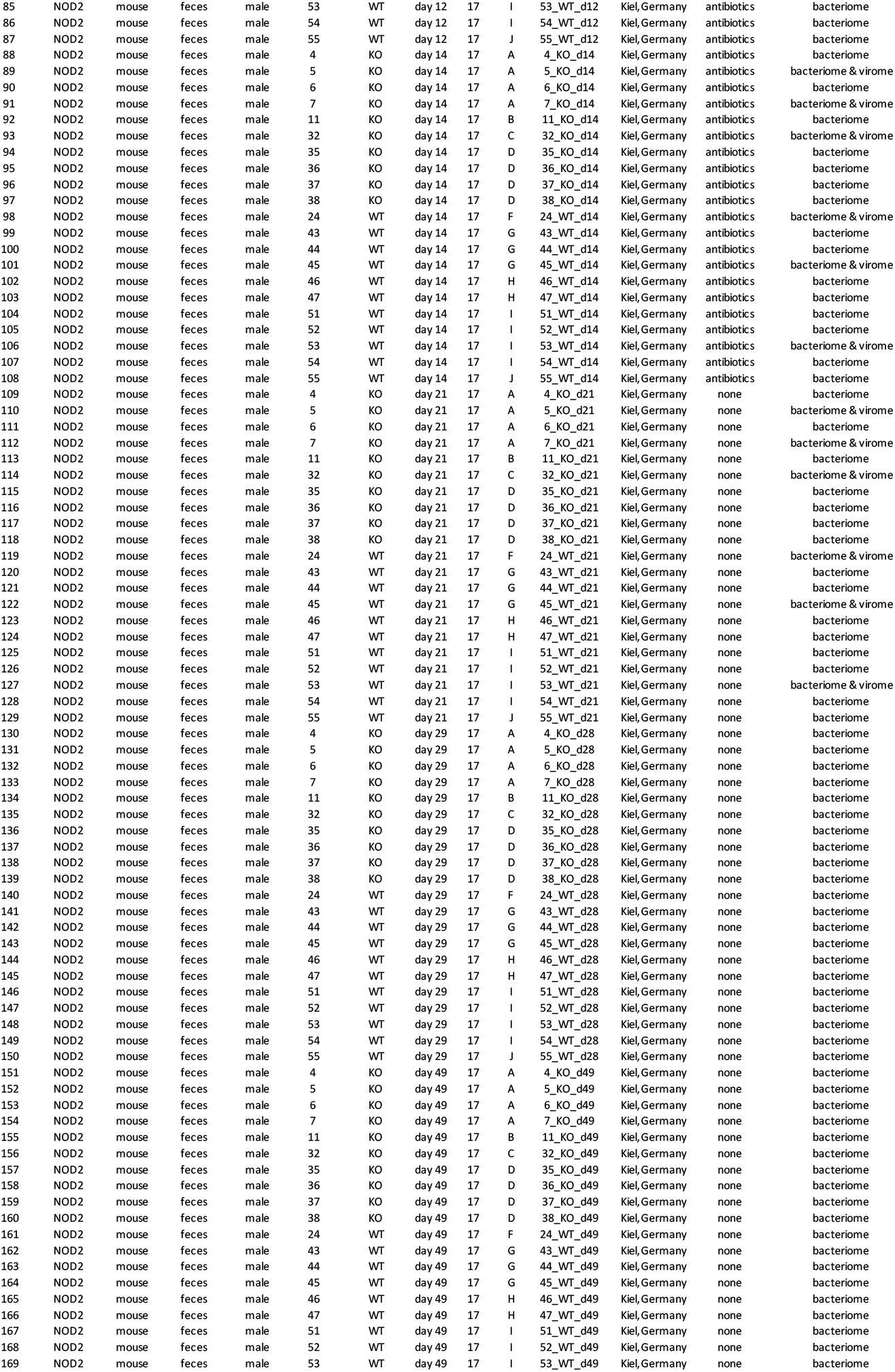
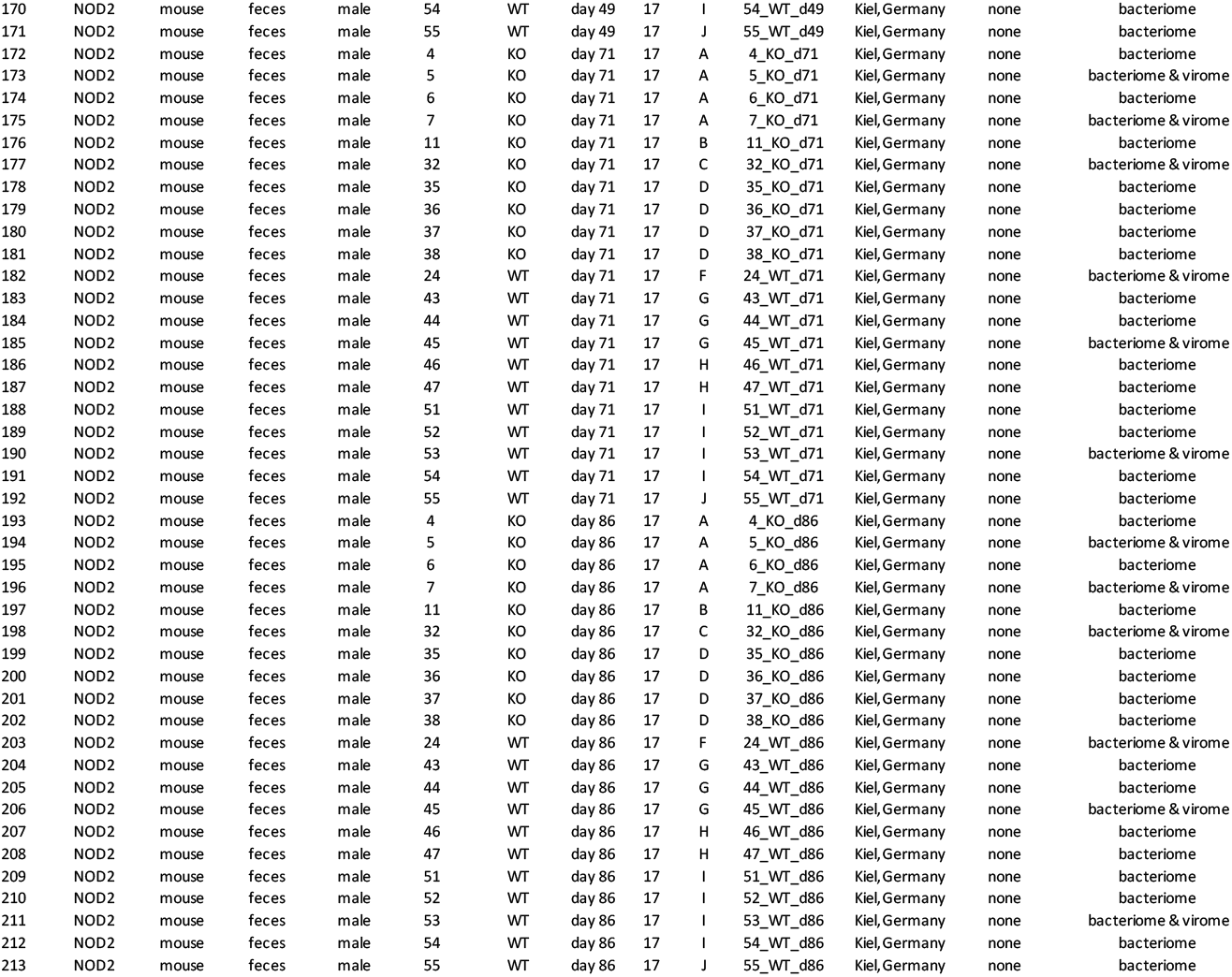

**Table S2.**
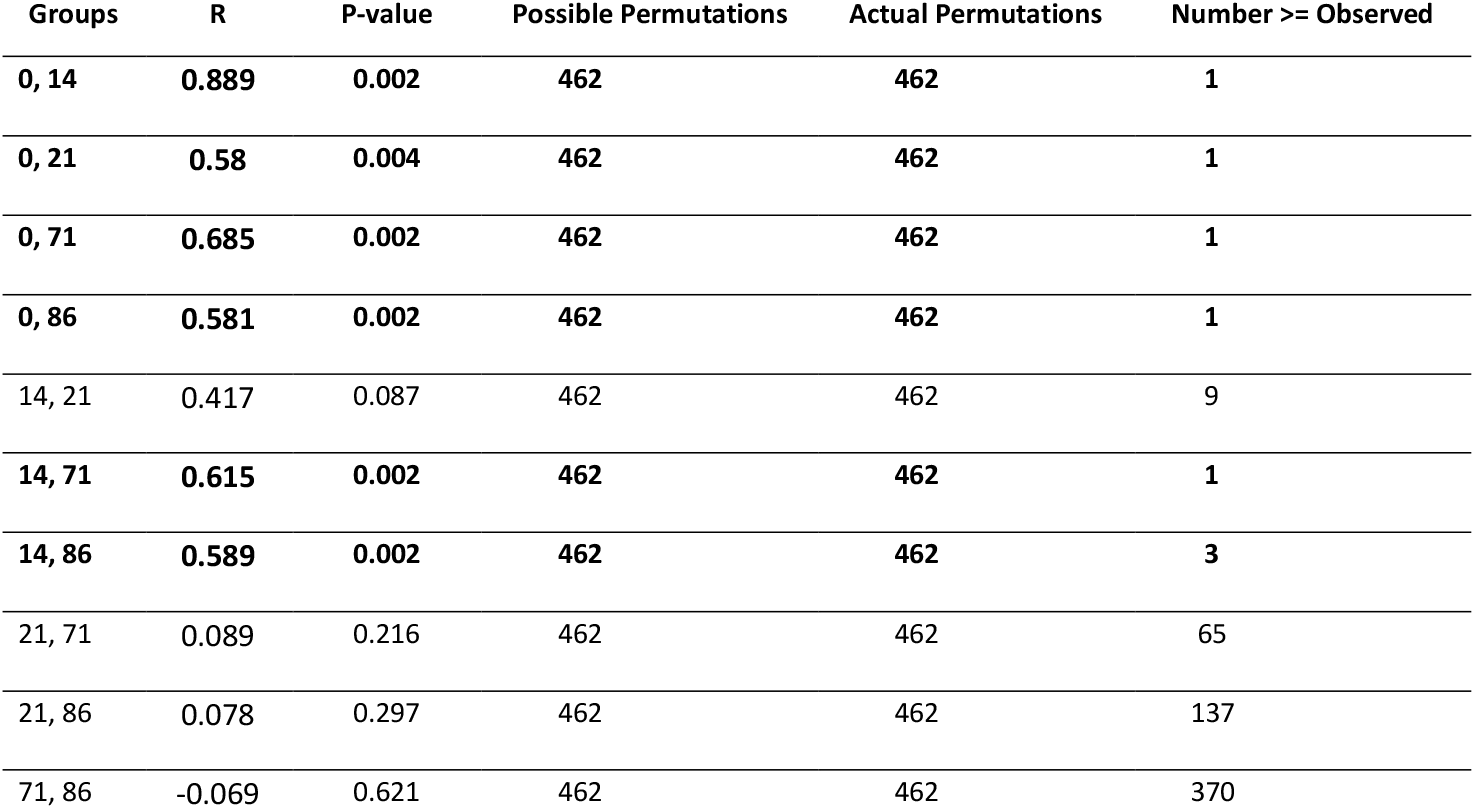
ANOSIM pairwise comparison of viral communities between time points. Day 0 before antibiotic perturbation; day 14 to day 86 post antibiotic perturbation.

**Table S3.**
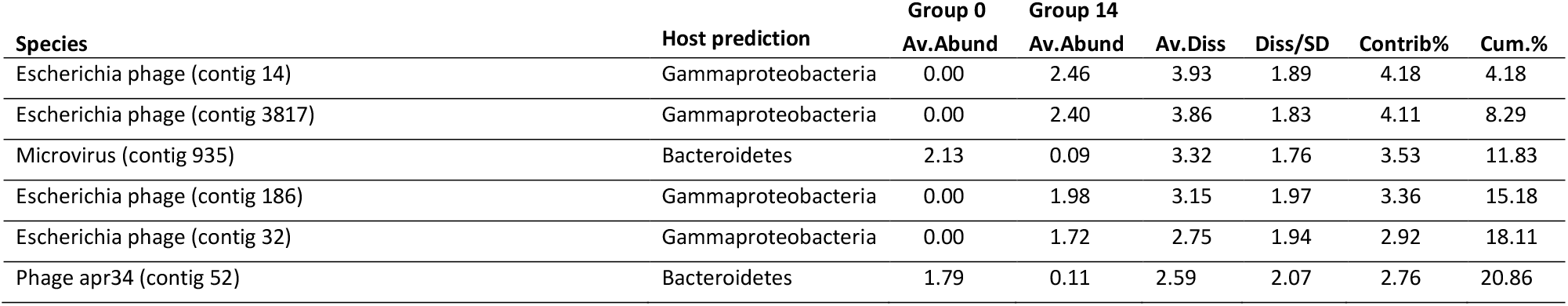
SIMPER analysis: pairwise test of differences between time points of groups (day 0 and day 14). Average dissimilarity = 93.92

**Table S4.**
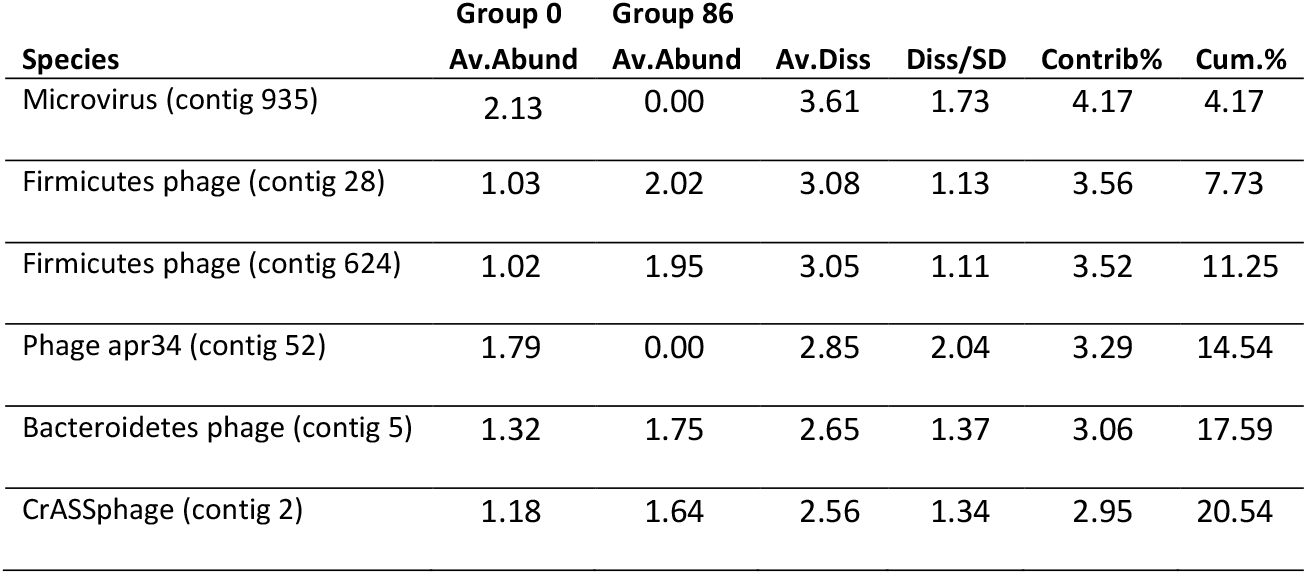
SIMPER analysis: pairwise test of differences between time points of groups (day 0 and day 86). Average dissimilarity = 86.64

**Table S5.**
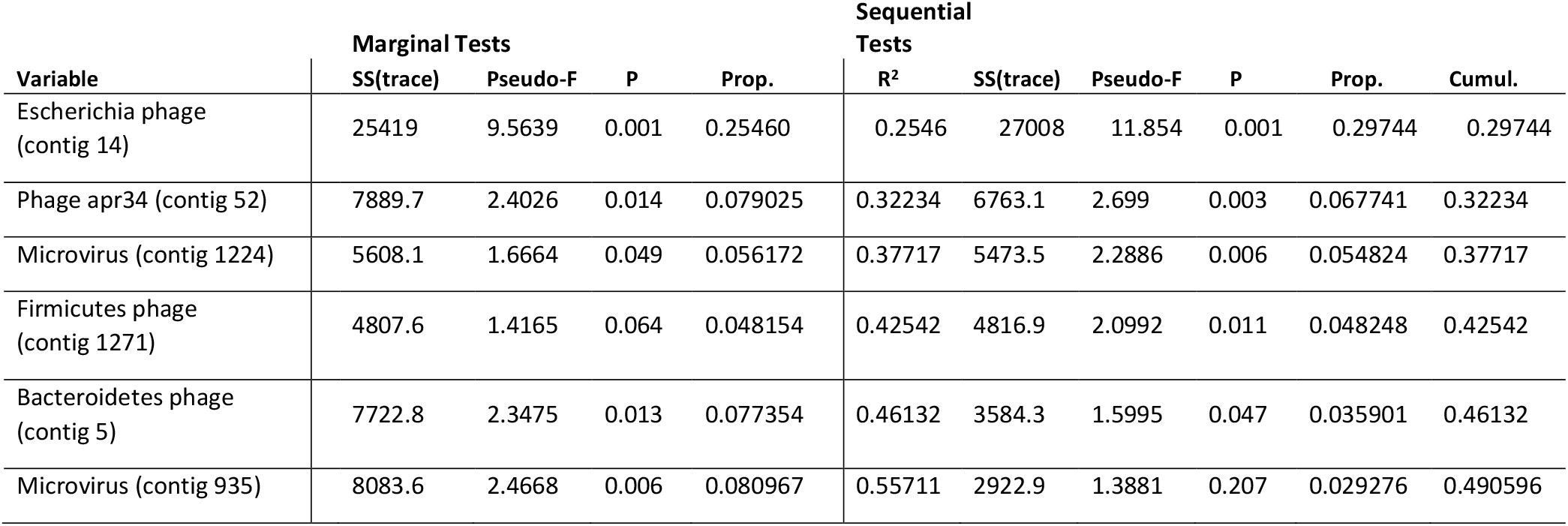
Proportion of variance in bacterial communities explained by the 29 most important viral OTUs as predictor variables in DistLM tests. Predictor variables were identified using adjusted R^2^ selection criteria and forward selection.

